# Single cell variation and rapid emergence of phenotypic heterogeneity in cell lines- a cautionary tale for devotees of CRISPR-Cas9

**DOI:** 10.1101/2024.09.27.615490

**Authors:** Hayley Bradley, John N Barr, Martin Stacey

## Abstract

The use and publication of research utilising CRISPR/cas9 in gene editing and knock outs (KOs) within cell lines is now widespread and has proved extremely powerful in the interrogation of gene function. However, the potential of experimental artefacts resulting from the need to generate single cell clones post gene-manipulation has been somewhat overlooked. In this study, we show that the commonly used pulmonary cell line A549 displays significant heterogeneity in terms of their gene expression, and that individual cells from a population exhibit dramatically different susceptibility to viral infection from a range of viruses including respiratory syncytial virus (RSV), influenza A virus (IAV), Hazara virus (HAZV), lymphocytic choriomeningitis virus (LCMV) and Bunyamwera virus (BUNV). Moreover, we demonstrate the rapid re-emergence of phenotypic heterogeneity even after cloning. These results demonstrate the need for caution in interpreting results from CRISPR screens and CRISPR-KO validation studies, especially in the study of viral infection.

## Introduction

The advent of CRISPR/Cas9 technology has revolutionised the area of molecular and cellular biology. It allows the rapid and inexpensive generation of genome-wide knockouts and gain-of-function libraries within cell lines in order to screen and interrogate gene function in multiple fields. Within the field of virology multiple studies have used CRISPR/cas9 technology to generate libraries of various cell lines to screen for host components essential for viral entry and replication. Moreover the use of CRISPR KO clones is increasingly used to validate the role of candidate genes, identified through other high throughput screens, in the viral life cycle. Although the use of somatic cell lines have obvious benefits there are potential pitfalls in using them in the study of virus entry and replication. Tissue culture adaption by viruses leading to the identification of potential erroneous receptors or host gene involvement in the viral life cycle have has been well documented. However, as CRISPR-KO within cell lines is becoming the go-to approach for the validation of gene function, this study provides a cautionary tale of its use in investigating host-viral interactions.

## Materials and Methods

### Cell culture and generation of clonal cell lines

A549 cells (ATCC, CCL-185) were maintained in high-glucose Dulbecco’s modified Eagle medium (DMEM; Sigma-Aldrich), supplemented with 10 % heat-inactivated foetal bovine serum (FBS), 100 μg of streptomycin/mL and 100 U of penicillin/mL. Cells were incubated in a humidified incubator at 37 °C with 5% CO_2_. The parental A549 cell population were seeded at a cell density of 1 cell/well into a 96-well cell culture plate and continuously monitored for single colony formation via both light microscopy and whole well scans using IncuCyte S3. Only wells which contained a single colony of cells were expanded, creating A549 clonal cell lines numbered 1-27. Clonal cell lines 7, 16, 24 and 26 were further passaged a minimum of 10 times and this process was then repeated, producing sub-clonal cell lines.

### Generation of CRISPR cell lines

Gene specific guide RNAs were designed using the CRISPick tool (Sanson, K. et al (2018). and subsequently cloned into the lentiCRISPR-V2 plasmid, a gift from Feng Zhang (Addgene plasmid # 52961; http://n2t.net/addgene:52961; RRID:Addgene_52961),

### Lentivirus production

Lentiviruses were produced containing the lentiCRISPR v2 plasmid with gDNA insert. HEK 293T cells were seeded in a 12-well plate at 5×10^5^ cells/well and transfected with the 0.4 μg lentiCRISPR v2, 0.3 μg pMD2.G (Addgene, 1225), and 0.3 μg psPAX2 (Addgene, 12260) using Jet Optimus transfection reagent (Polyplus). 48 hours post transfection the supernatant was harvested and clarified via centrifugation at 500xg, 5 minutes followed by 0.45 μm filtration. Lentivirus-containing medium was used immediately, or stored at 4°C.

### Knockout generation

Gene KOs were generated in A549 cells through infection with lentiviruses containing the lentiCRISPR v2 plasmid with respective gDNA inserts. A549 cells were seeded at 1×10^5^ cells/well in a 24-well plate, incubated overnight, and infected with the lentivirus-containing supernatant of transfected 293Ts supplemented with 8 μg/mL Polybrene. At 48 hours post-infection (hpi), media was replaced with 10% FBS DMEM containing 1 μg/mL puromycin. Cells were incubated for a further 7 days to eliminate uninfected cells, before cloning as described above.

### Knockout validation

Genomic DNA was isolated from 0.5 × 10^6^ of both wild type (WT) and KO A549 cells using the Qiagen ‘Flexigene DNA Kit’ protocol (Qiagen, 2023). PCR primers were designed for KO regions and used to amplify these regions in both WT DNA and KO DNA. PCR success was validated using 1.5% agarose gel electrophoresis and PCR products were sequenced. The online ‘ICE CRISPR Analysis Tool’ from Synthego was used to analyse and compare knockout sequences of different target genes against their respective WT counterparts.

### Virus strains and generation of recombinant reporter viruses

GFP-PA (polymerase acidic protein) labelled IAV (H1N1 WSN/33 strain) was rescued as described previously (PMID: 10430945). GFP-expressing RSV in which the *gfp* gene had been inserted as an independent transcriptional unit in the first position in the HRSV gene order was purchased from ViraTree (RSV-GFP1).

The generation of recombinant, eGFP-expressing Hazara virus and lymphocytic choriomeningitis virus (rHAZV-eGFP/rLCMV-eGFP) were described previously by (PMID: 32581103 PMID: 38874413, respectively). Using this established protocol, recombinant Bunyamwera virus harbouring an eGFP reporter gene was generated (rBUNV-eGFP). In all cases, the eGFP open reading frame (ORF) was inserted into plasmids encoding the S-segment of each virus, upstream of the NP ORF, and was linked by a self-cleaving P2A region. This resulted in the generation of recombinant reporter viruses which express eGFP alongside viral protein synthesis.

### Virus infection

A549 clones or subclones were seeded at a density of 1×10^4^ cells/well in a 96-well plate and left to adhere and grow overnight. Culture media was removed and cells were washed in 1x phosphate buffered saline (PBS), before infection with virus in low-serum DMEM. All infections were performed at MOI 0.1, and left for either 24 hrs (rBUNV-eGFP, rHAZV-eGFP, rLCMV-eGFP, RSV-GFP) or 48 hrs (IAV-PA-GFP). Infections and mock infections were performed in triplicate and the averages were taken. Infection was analysed via the quantification of GFP-expressing cells using IncuCyte S3.

### RNA extraction and analysis

3 biological replicates from A549 clonal cell lines 1, 16 and 26 were collected and RNA was extracted using RNeasy kit (Qiagen). Standard NGS RNA-seq using PolyA selected RNA 150 bp pair-end reads was performed by Genewiz. Raw FASTQ files were processed and analysed using the Galaxy Platform (https://usegalaxy.org/). Raw RNA-seq paired-end reads were subjected to quality control using FastQC, low-quality bases and adapter sequences were removed using Cutadapt, subsequent trimmed reads were aligned to the reference genome (GRCh38) using HISAT2, and differential gene expression analysis was conducted using DESeq2 in R (v4.0.3). Raw read counts were normalized, and differential expression was determined using a negative binomial generalized linear model. Genes with an adjusted p-value ≥0.05 were considered significantly differentially expressed (PMID: 33835453).

### Statistical Analysis

The statistical significance of data were determined by performing one-way ANOVA analysis, with a p-value of ≤0.05 considered significant.

## Results

### Individual CRISPR KO clones display highly variable phenotypes despite identical gene disruption

Previous studies have demonstrated the critical nature of the endosomal environment in the entry and fusion of enveloped viruses, especially those of the Peribunyaviridae family (PMID: 26677217, PMID: 37735161). Therefore, in order to identify potential genes involved, an siRNA based screen of selected ion channels was performed on the prototypic member, Bunyamwera. Amongst promising candidate genes in the commonly used A459 cell line that reduced viral entry was the potassium (K^+^) inwardly-rectifying channel, subfamily J, member 13 (*KCNJ13*), which encodes the K_ir_7.1 channel. To circumvent potential issues with the deficiencies of transfection efficiency, and incomplete protein ablation via transient siRNA, we chose to validate this hit using a lentiviral CRIPSR-KO strategy. Following disruption of the *KCNJ13* gene, the subsequent analysis of the *KCNJ13* gene amplicons via ICE-Sythnego software (https://ice.synthego.com/) demonstrated that initial transduction of gene-specific guide-RNA lead to a population of mixed and incomplete *KCNJ13* gene disruptions (gene disruption and knockout scores of 64% and 59%, respectively). Therefore, to generate homozygous complete KOs for subsequent studies, single cell clones were isolated and sequence verified (Fig 1A, B). These clones, along with the parental A549 cell population, were infected with BUNV-eGFP at MOI 0.1 for 24 hours (Fig 1B). Worryingly, despite the identical nature of the gene disruption in all 3 clones (a 10bp frameshift-associated deletion, Fig 1A), they showed significant variability with each other and with the parental population of A549 cells, ranging from 69%-120% levels of viral infection. This indicated that there could be inherent variability in the infectibility of individual A549 cell clones with BUNV.

**Figure 1.**
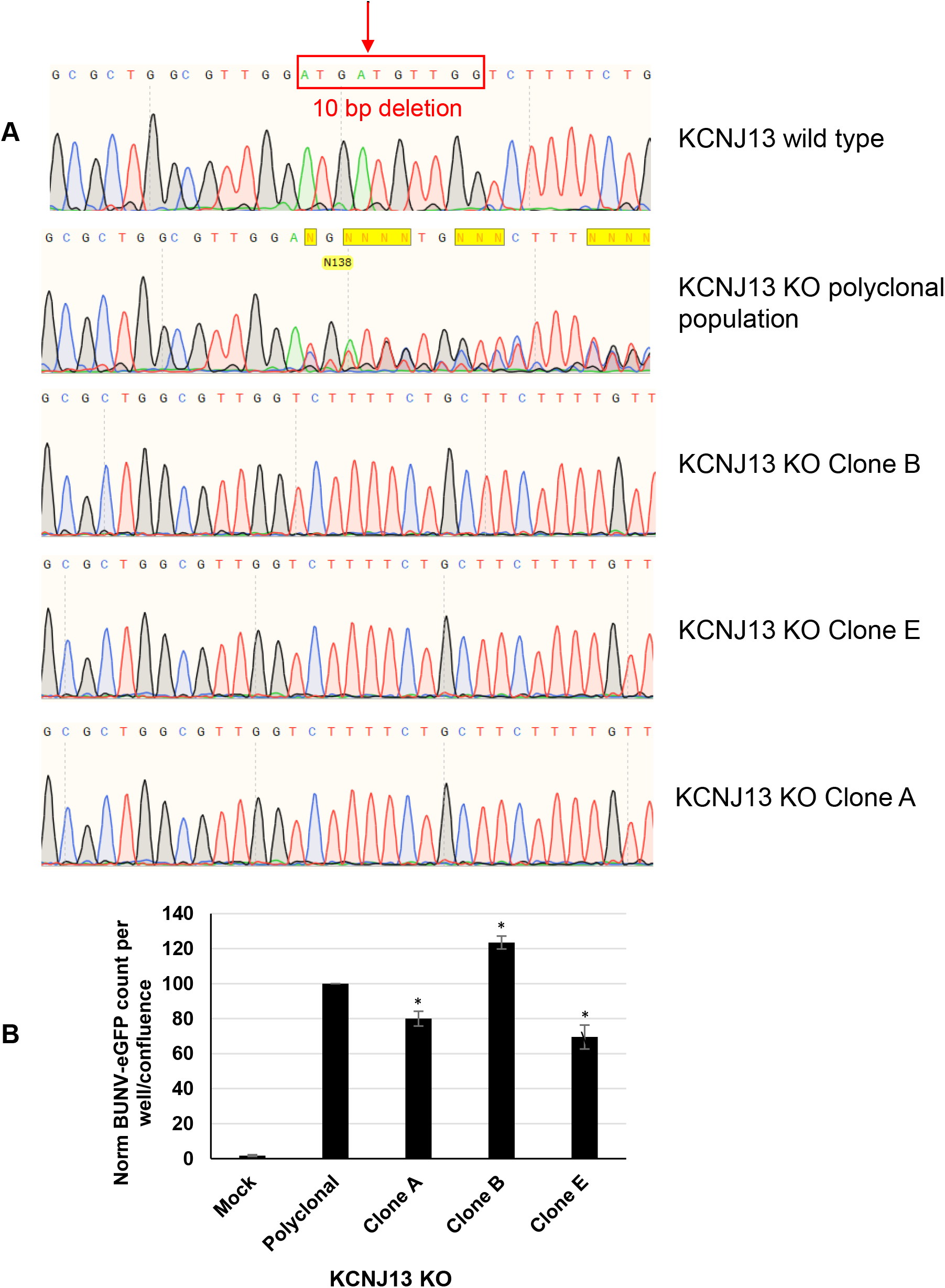
A459 KCNJ13 KO clones show variable susceptibility to BUNV infection. **A)**. Genome sequence traces of KCNJ13 gene targeted with specific CRISPR-Cas9 guide-RNA. Red arrow denotes predicted Cas9 cleavage site, box shows 10bp deletion found in clones A, B and E. Genomic sequence trace of polyclonal population is shown displaying overlapping sequences due to multiple different sequences generated by non-homologous end joining post-Cas9 mediated DSB. ICE-Synthego Analysis predicts gene disruption and knockout scores of 64% and 59%, respectively. **B)** Clonal cell lines were infected or mock-infected with BUNV-eGFP at MOI 0.1 for 24 hours and infection was assessed via quantification of eGFP-expressing cells by IncuCyte S3. Experiments were performed in triplicate and average of n=4 repeats is shown, error bars represent SE. Values are normalised to the parental cell line. *p-value <0.05

### Individual A549 cell clones display significant variability to infection by a range of viruses

In order to explore whether the variability of infection was due to process of CRISPR manipulation or phenotypic heterogeneity within the original parental cell line, numerous clones were generated via clonal dilution and expansion, and subsequently infected with rBUNV-eGFP (Fig 2A, B). As with the CRISPR-KO generated clones, despite low intra-experimental variability, susceptibility to infection by the WT-derived clones varied significantly, ranging from ∼50% to 120% when compared to the mixed parental population (Fig 2B). To test whether this was limited to BUNV or a broader phenomenon, the clones were infected with other members of the *Bunyaviricetes* class; rHAZV-eGFP and rLCMV-eGFP. Similarly, the WT A549 clones showed significant and variable susceptibility to infection by these additional Bunyaviruses (Fig. 2 C, D). To further investigate whether this phenomenon was specific to Bunyaviruses or could be applied to other viruses, we next infected the WT A549 clones with two respiratory, negative-sense RNA viruses; RSV and IAV (Fig 2E-F). IAV in particular displayed a dramatic variability of infection of clones, between 20-120% when compared to the mixed parental population. Of note, individual clones (eg 26) varied significantly between viruses showing 20% infection with IAV and 120% infection with RSV. This suggests that clonal susceptibility is unlikely to be due to a single factor that would affect multiple virus types such as cell viability, global translational/endocytic capacity or levels of the anti-viral response.

**Figure 2.**
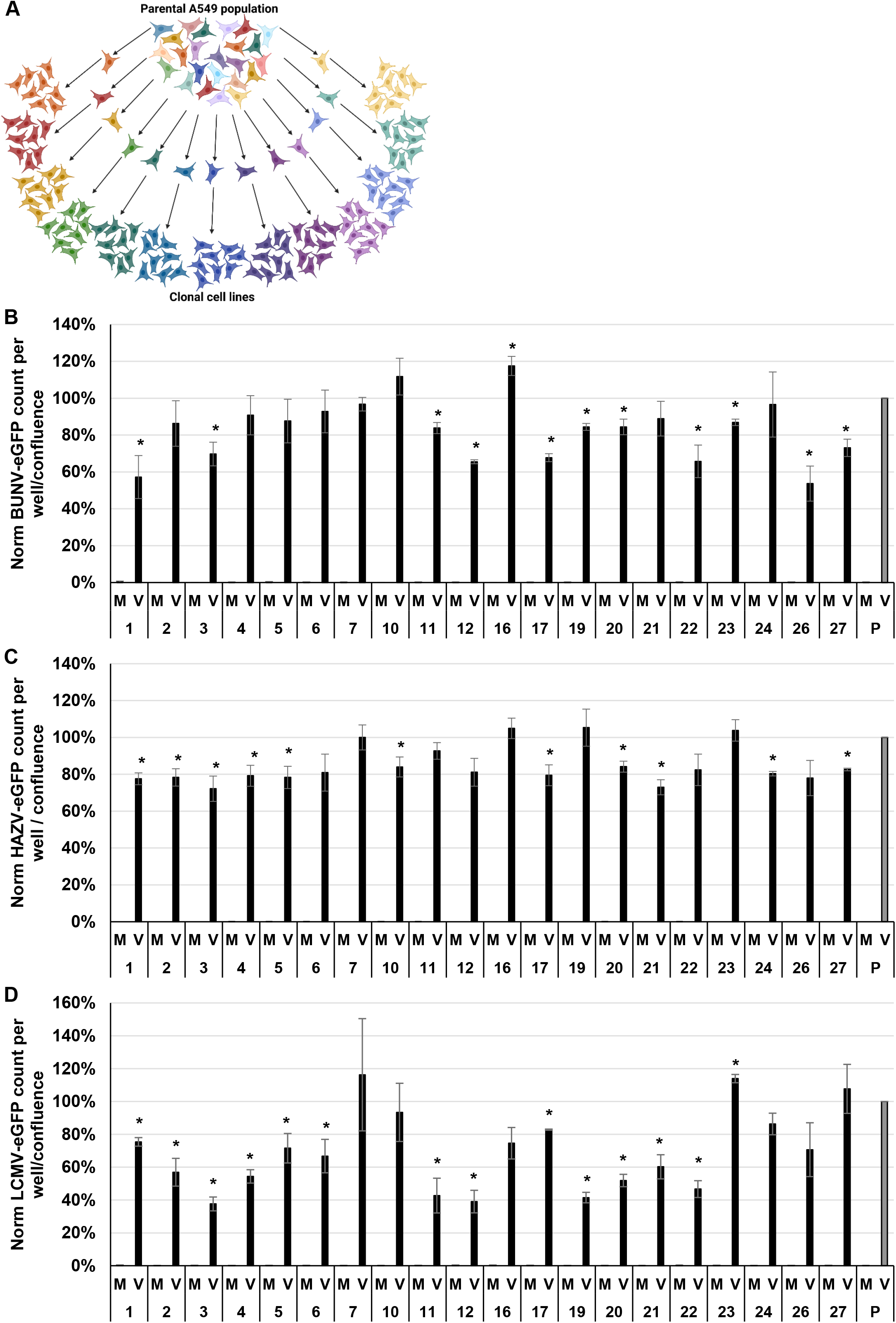
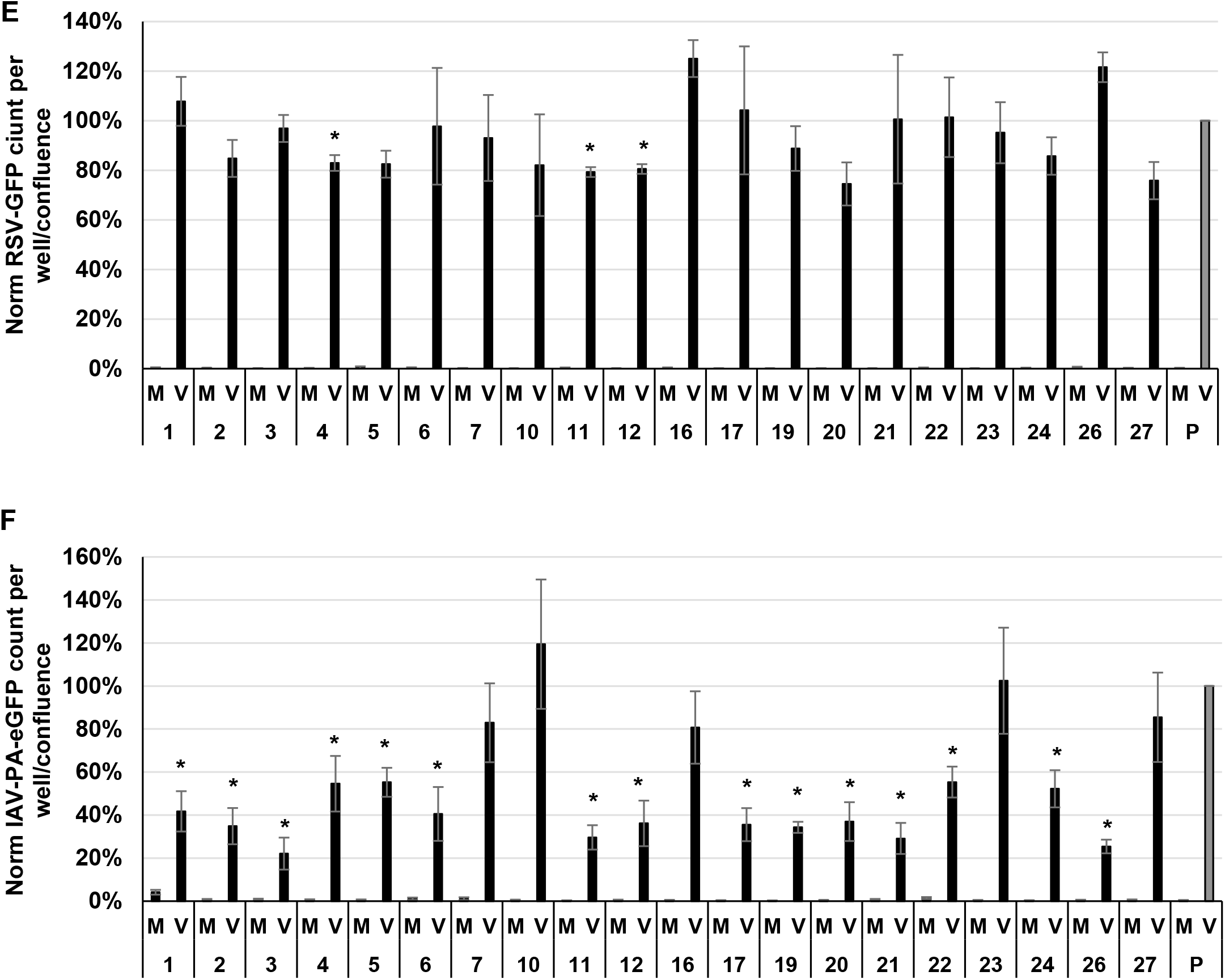
A549 clones of a parental cell line have variable susceptibilities to Bunyavirus infection. **A**. The parental A549 cell population were seeded into a 96-well plate at 1 cell per well. Cells were monitored for colony formation and single colonies were expanded to create clonal cell lines 1-27. **B-D**. Clonal cell lines were infected or mock-infected with BUNV-eGFP, HAZV-eGFP or LCMV-eGFP at MOI 0.1 for 24 hours and infection was assessed via quantification of eGFP-expressing cells by IncuCyte S3. Experiments were performed in triplicate and the average of n=3 is shown, error bars represent SE. Values are normalised to the parental cell line. *p-value <0.05.

### Phenotypic heterogeneity of A549 clones rapidly re-emerges after passage

To circumvent potential experimental artefacts due to clonal heterogeneity, a potential strategy would be to initiate CRISPR experiments using single clones. However, this approach would only prove suitable if clones maintained their phenotypic homogeneity. To investigate this, selected WT A549 clones (7, 16, 24, 26) were passaged a further 10 times and re-cloned (Fig 3A). The new subclones were then infected with rBUNV-eGFP. After only 10 passages, heterogeneity in the susceptibility to infection had returned to all clones (Fig 4B-E). Some subclones displaying only 20% infection compared to others derived from the same initial clone (e.g. subclones 26.1 and 26.6, Fig 4E). Furthermore, the average phenotype of the each of the sets of subclones did not retain the phenotype their parental clone, i.e. highly susceptible/resistant clones on average did not continue to be highly susceptible/resistant (Supplementary Fig. 1), suggesting the rapid and stochastic re-emergence of heterogeneity upon cell passage. From this, we conclude that performing CRISPR experiments on single A549 clones would not resolve the issue that we initially described regarding the highly variable phenotypes in terms of viral infection.

**Figure 3.**
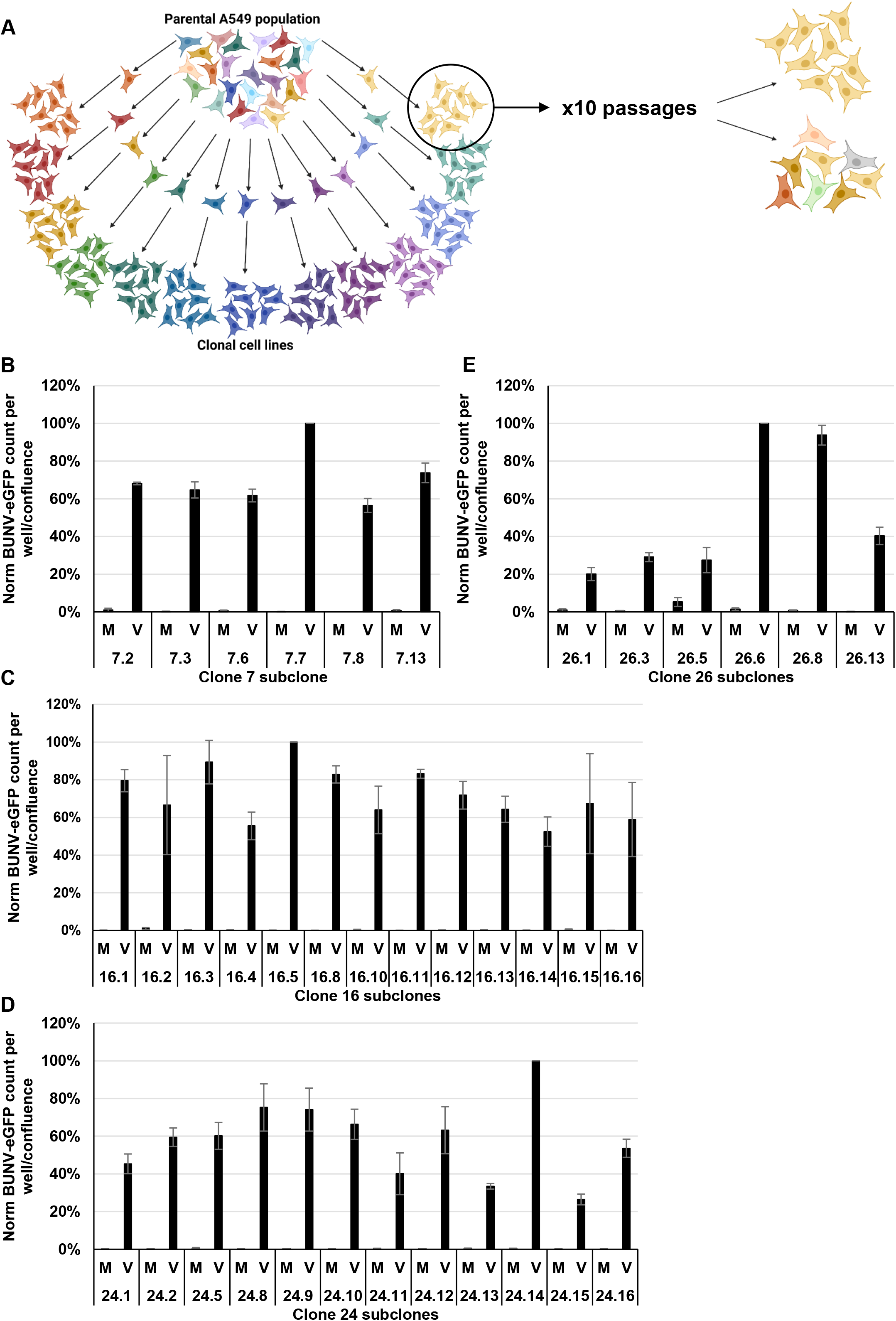
A549 clones of a parental cell line have variable susceptibilities to respiratory virus infection. **A**. Clonal cell lines were infected or mock-infected with RSV-GFP at MOI 0.1 for 24 hours and infection was assessed via quantification of eGFP-expressing cells by IncuCyte S3. **B**. Clonal cell lines were infected with IAV-PA-GFP at MOI 0.1 for 48 hours and infection was assessed as in A. Experiments were performed in triplicate and average of n=2 and n=3 is shown for A and B respectively. Error bars represent SE. Values are normalised to the parental cell line. *p-value <0.05. **Clonal A549 cell lines do not retain their homogeneity following passage. A**. Selected clonal A549 cell lines were passaged 10 times in cell culture and tested to see if they retained their clonality or returned into a heterogeneous population in terms of their susceptibility to virus infection. **B-E**. Subclones generated from A549 clonal cell lines 7, 16, 24 and 26 were infected or mock-infected with BUNV-eGFP at MOI 0.1 for 24 hours. Infection was assessed by quantifying eGFP expression by IncyCyte. Experiments were performed in triplicate and n=2/3, error bars represent SE. *p-value<0.05.

**Figure 4.**
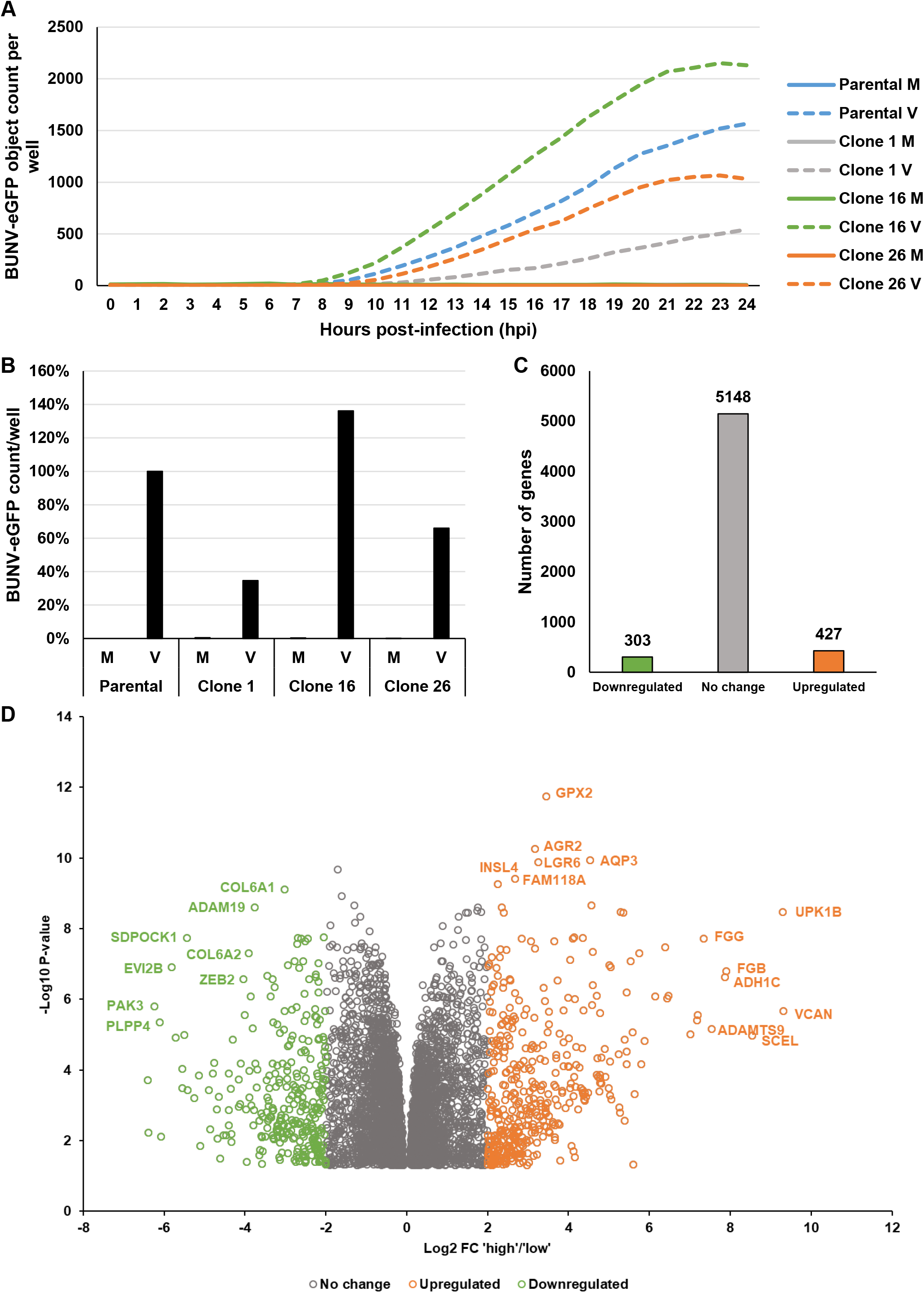
Clonal A549 cell lines exhibiting the highest or lowest susceptibility display significant changes in gene expression. **A**. Clonal A549 cell line 16 (higher susceptibility to BUNV-eGFP infection) and lines 1 and 26 (lower susceptibility to BUNV-eGFP infection) were infected or mock-infected with BUNV-eGFP at MOI 0.1. eGFP expression was assessed hourly for 24 hours and the parental cell line was included for comparison. A direct comparison of this infection after 24 hours is shown in **B**. Infection was performed in triplicate in a 96 wp, n=1. C-D. RNA-seq analysis of

### A549 clones exhibit differential RNA-expression profiles

To investigate the potential genes or set of genes responsible for the differences in clonal susceptibility to virus infection, the RNA-expression profiles of different WT A549 clones were analysed. Focussing specifically on BUNV, clones exhibiting the highest (16) or lowest (1 and 26) susceptibility to infection were chosen for transcriptomic analysis. Cells taken from the same flask at the same time (i.e same passage number) were used to perform virus infections as well as being the source of RNA. This was to ensure that we had an accurate picture of the phenotype of the cells that were subjected to RNA-seq. BUNV-eGFP expression was analysed every hour for 24 hours. Clone 16 (highly infectible) showed the earliest and highest eGFP expression over the full time course, with the more resistant clones (1 and 26) exhibiting later and less eGFP expression than the parental A549 cell population control (Fig 4A,B). This corroborated the phenotypes observed for these clonal lines in the previous experiments.

Despite being obtained from the same parental cells, the transcriptional profile between high and low clones differed greatly with over 400 genes expressed at significantly higher levels (>log2 fold increase) in the highly susceptible cells whereas >300 genes were expressed at high level in the low susceptible cell. Gene Ontology/KEGG pathway analysis yielded a variety of significantly enriched terms in both sets of clones including protein glycosylation, cell-cell interactions and immune receptor activation; all of which could have the potential to either facilitate/reduce viral infection.

## Discussion

The use of CRISPR-Cas9 technology is now commonplace in molecular virology as well as other fields. It has been successfully used in genome-wide knockout screens to identify novel virus receptors and key host components essential in the virus life cycle (PMID: 32522852, PMID: 35086559). Its use in convenient tractable cell lines is also often employed to verify/validate initial gain-of-function /loss-of-function screens as well as the role of candidate genes from multiple other experimental approaches. One such cell line is the primary lung epithelial cell line, A549 that is commonly used in the studies of respiratory and other viruses including SARS-CoV2, RSV, IAV and Bunyaviruses (PMID: 35891350, PMID: 33109690, PMC10949467), as well in the study of lung pathology (PMID: 30505938). A459 cells are considered stable, immunocompetent and retain, at least in part, the phenotype of alveolar Type II pulmonary epithelial cells from which they are derived. In this study, we show that the heterogeneity within A459 cells has the potential to generate erroneous results and experimental artefacts. When compared, individual cloned cells displayed a large number of differentially expressed genes and dramatic phenotypic differences in terms of their susceptibly to viral infection. This may well be likely in other types of phenotype screens and in other cell lines. Moreover, even after cloning, individual clones rapidly (10 passages) developed significant heterogeneity similar to that of the parental cell line. This prevents a strategy whereby initial cloning of a parental cell line would circumvent the issue of phenotypic heterogeneity. In summary, these results highlight the need for caution when using genetically manipulated cloned cell lines in the validation of gain/loss-of function experiments, and the suggest the use of polyclonal population and other complementary experimental approaches.

## Acknowledgements

We would like to thank Dr Amelia Shaw, Dr Eleanor Todd and Sophia Qias (Dr John Barr’s lab, University of Leeds) for the gifts of rLCMV-eGFP, IAV-PA-GFP and rHAZV-eGFP, respectively. We acknowledge funding from BBSRC grant Grant BB/V007467/1(HP).

---

**Supplementary figure 1.**
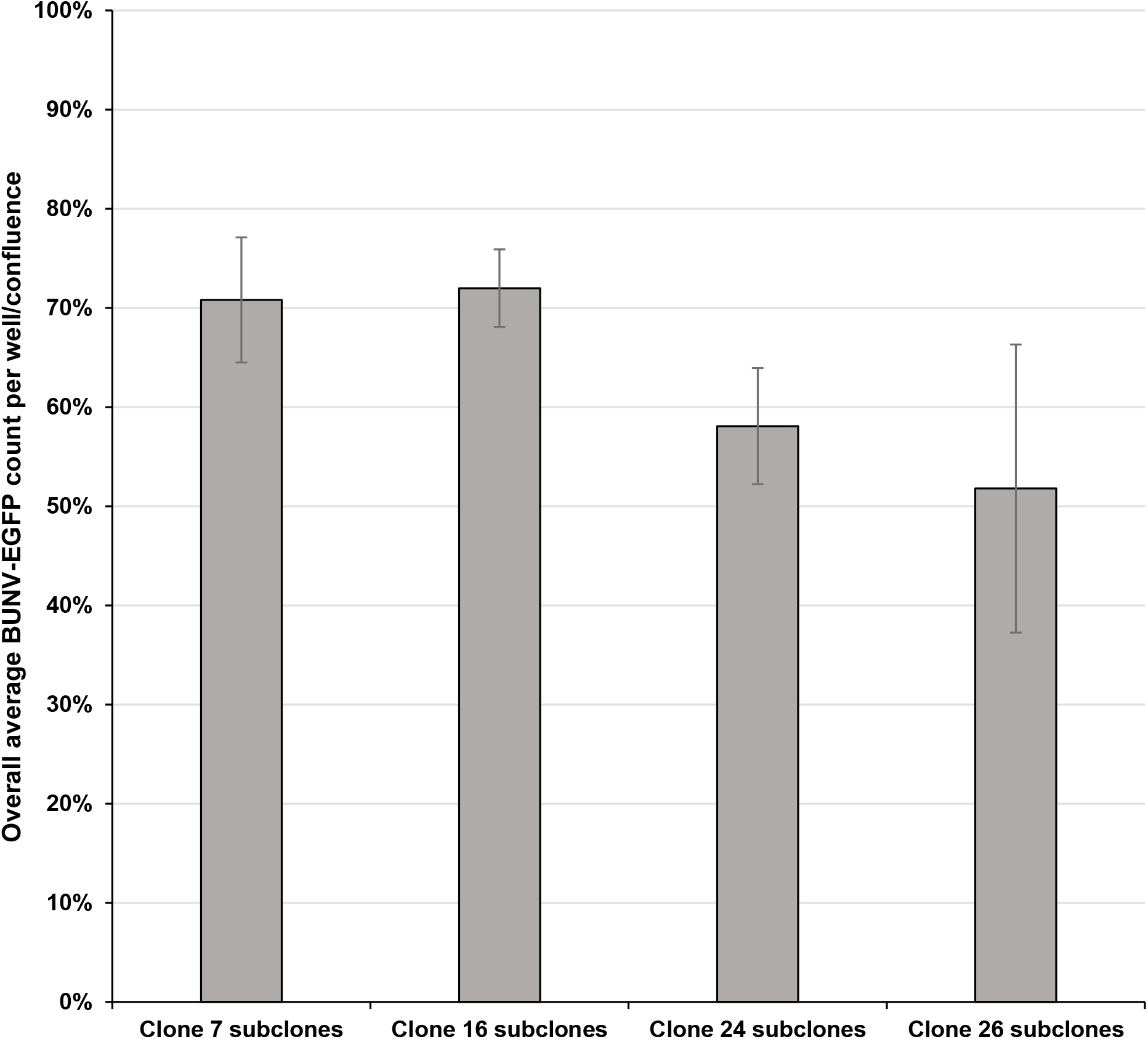
After 10 passages subclones do not retain their parental phenotype. Multi subclones of clone 7, 16, 24, and 26 (between 6-13) were infected with BUNV-eGFP, at an of MOI 0.1 for 24 hours. Infection was assessed via quantification of eGFP-expressing cells by IncuCyte S3. Within individual experiments infection were performed in triplicate and the average of 3 independent experiments is shown, error bars represent SE. *p-value <0.05

